# *MAU2* and *NIPBL* variants in Cornelia de Lange syndrome reveal MAU2-independent loading of cohesin and uncover a protective mechanism against early truncating mutations in NIPBL

**DOI:** 10.1101/477752

**Authors:** Ilaria Parenti, Farah Diab, Sara Ruiz Gil, Eskeatnaf Mulugeta, Valentina Casa, Riccardo Berutti, Rutger W.W. Brouwer, Valerie Dupé, Juliane Eckhold, Elisabeth Graf, Beatriz Puisac, Feliciano Ramos, Thomas Schwarzmayr, Thomas van Staveren, Wilfred F. J. van IJcken, Tim M. Strom, Juan Pié, Erwan Watrin, Frank J. Kaiser, Kerstin S. Wendt

## Abstract

Cornelia de Lange syndrome (CdLS) is a rare developmental disorder caused by mutations in genes related to the cohesin complex. For its association with chromatin, cohesin depends on a heterodimer formed by NIPBL and MAU2, which interact via their respective N-termini. Variants in *NIPBL* are the main cause of CdLS and result in *NIPBL* haploinsufficiency.

Using CRISPR, we generated cells homozygous for an out-of-frame duplication in *NIPBL*. Remarkably, alternative translation initiation rescued NIPBL expression in these cells and produced an N-terminally truncated NIPBL that lacks MAU2-interaction domain, causing a dramatic reduction of MAU2 protein levels. Strikingly, this protective mechanism allows nearly normal amounts of cohesin to be loaded onto chromatin in a manner that is independent of functional NIPBL/MAU2 complexes and therefore in contrast to previous findings.

We also report the first pathogenic variant in *MAU2*, a deletion of seven amino acids important for wrapping the N-terminus of NIPBL within MAU2. The mutation causes dramatic reduction of MAU2 heterodimerization with NIPBL, hence undermining the stability of both proteins.

Our data confirm *NIPBL* haploinsufficiency as the major pathogenic mechanism of CdLS and give new insights into the molecular mechanisms responsible for this neurodevelopmental disorder. Our work also unveils an alternative translation-based mechanism that protects cells from out-of-frame variants of NIPBL and that may be of relevance in other genetic conditions.

## INTRODUCTION

Cohesin is a highly conserved protein complex essential for cell survival. In humans, the complex is composed of three structural core subunits named SMC1A, SMC3 and RAD21, which together form a ring-shaped structure that topologically encircles DNA^1,2^. This ability allows cohesin to carry out a large spectrum of chromatin-related functions, including sister chromatid cohesion, DNA repair, transcriptional regulation and three-dimensional organization of chromatin^3^. In order to accomplish these essential tasks, cohesin needs to interact with chromatin. Cohesin’s binding to DNA depends on the heterodimer formed by NIPBL and MAU2, also known as cohesin loader or kollerin complex^4,5^. Importantly, NIPBL is also known to play a role in transcriptional regulation independently of cohesin^6^. The interaction with cohesin is mediated by the C-terminus of NIPBL^7^ and mutations affecting this domain result in reduced interaction between NIPBL and RAD21^8^. The interaction between NIPBL and MAU2 is instead mediated by their respective N-termini; precisely, the first 38 amino acids of NIPBL interact with amino acids 32-71 of MAU2^9^. Recent evidence suggested that MAU2 is required for the correct folding of the N-terminus of NIPBL and that NIPBL is unstable in the absence of MAU2^7,10^. RNA interference experiments similarly demonstrated that depletion of NIPBL greatly reduces the cellular levels of MAU2^5^, consistent with the notion that the physical association of these two proteins is required for their stability. Expression levels of cohesin and NIPBL are crucial for cells and tightly regulated. Mutations in different subunits of the cohesin complex (*SMC1A, SMC3, RAD21*) or of its regulators (*NIPBL, HDAC8*) are responsible for the onset of the multisystem neurodevelopmental disorder known as Cornelia de Lange syndrome (CdLS, OMIM #122470, 300590, 610759, 614701 and 300882)^11–16^. CdLS is characterized by pre- and post-natal growth retardation, intellectual disability, developmental delay, limb anomalies and distinctive facial features^17^.

The most common cause of CdLS is *NIPBL* haploinsufficiency, which is reported in approximately 65% of patients^18–20^, while up to 15% of cases are collectively accounted for by mutations in cohesin genes and additional transcriptional regulators or chromatin remodelers^13,21–27^. In contrast, pathogenic mutations in *MAU2* have never been reported, even though its function is essential for the stability of NIPBL. Of importance, a decrease as little as 15% of *NIPBL* expression was shown to cause distinct mild CdLS phenotypes. In addition, *NIPBL* transcript levels never fall below 60% in cell lines of patients with CdLS, suggesting that human development is extremely sensitive to *NIPBL* gene dosage^28,29^. Different types of heterozygous alterations have been described in association with *NIPBL*, including missense substitutions, nonsense and splicing mutations as well as out-of-frame deletions and insertions^18–20^. Notably, truncating mutations in *NIPBL* are mostly associated with a more severe phenotype and with a higher frequency of limb reductions or malformations in comparison to missense substitutions^20^. In apparent contradiction with this observation, some patients with very early truncations present with mild to moderate phenotypes and do not always display dramatic reductions of the upper limbs^18,30,31^, raising the interesting possibility that molecular consequences of *NIPBL* nonsense mutations depend on their position along the protein sequence.

In this work, we investigated the mechanisms responsible for the onset of such phenotypes. Genome editing of HEK293 cells provided a successful strategy to study the effects of homozygous early truncations in *NIPBL* on its binding partner MAU2 and on its substrate cohesin. Our results indicate that alternative translation start sites are used in the presence of early truncations, leading to the expression of an N-terminally truncated form of NIPBL that is able to mediate cohesin loading and does not depend on MAU2 for its stability. Accordingly, MAU2 levels are dramatically reduced in the edited cells.

We also identified the first pathogenic variant in *MAU2*, which provides further insight into the tightly regulated association between the two subunits of the kollerin complex. The mutation, an in-frame deletion of seven amino acids, impairs the interaction between NIPBL and MAU2, affecting the total level of both proteins.

Altogether, our data shed new light on the relationship between NIPBL and MAU2 and unveil an unprecedented implication of MAU2 in the pathogenesis of CdLS as well as a protective mechanism that dodges the deleterious effects of certain *NIPBL* mutations.

## RESULTS

### Cells with a homozygous loss-of-function mutation in *NIPBL* express an N-terminally truncated NIPBL and lack MAU2

To investigate the molecular mechanisms responsible for the pathogenesis of CdLS in the presence of early truncations in *NIPBL,* we performed CRISPR/Cas9 genome editing in HEK293 cells with sgRNAs targeting exon 2 of *NIPBL* with the purpose of generating an early truncating mutation through the error-prone repair mechanism non-homologous end joining. This experiment led to the establishment of two independent isolated clones carrying the same out-of-frame duplication of T (c.39dup) in a homozygous state, predicted to result in the null protein p.(Ala14Cysfs*5) (Figure 1a). Interestingly, despite the presence of a homozygous loss-of-function mutation in *NIPBL*, no difference between wild type and genome-edited cells was observed with regard to cell viability and proliferation.

**Figure 1.**
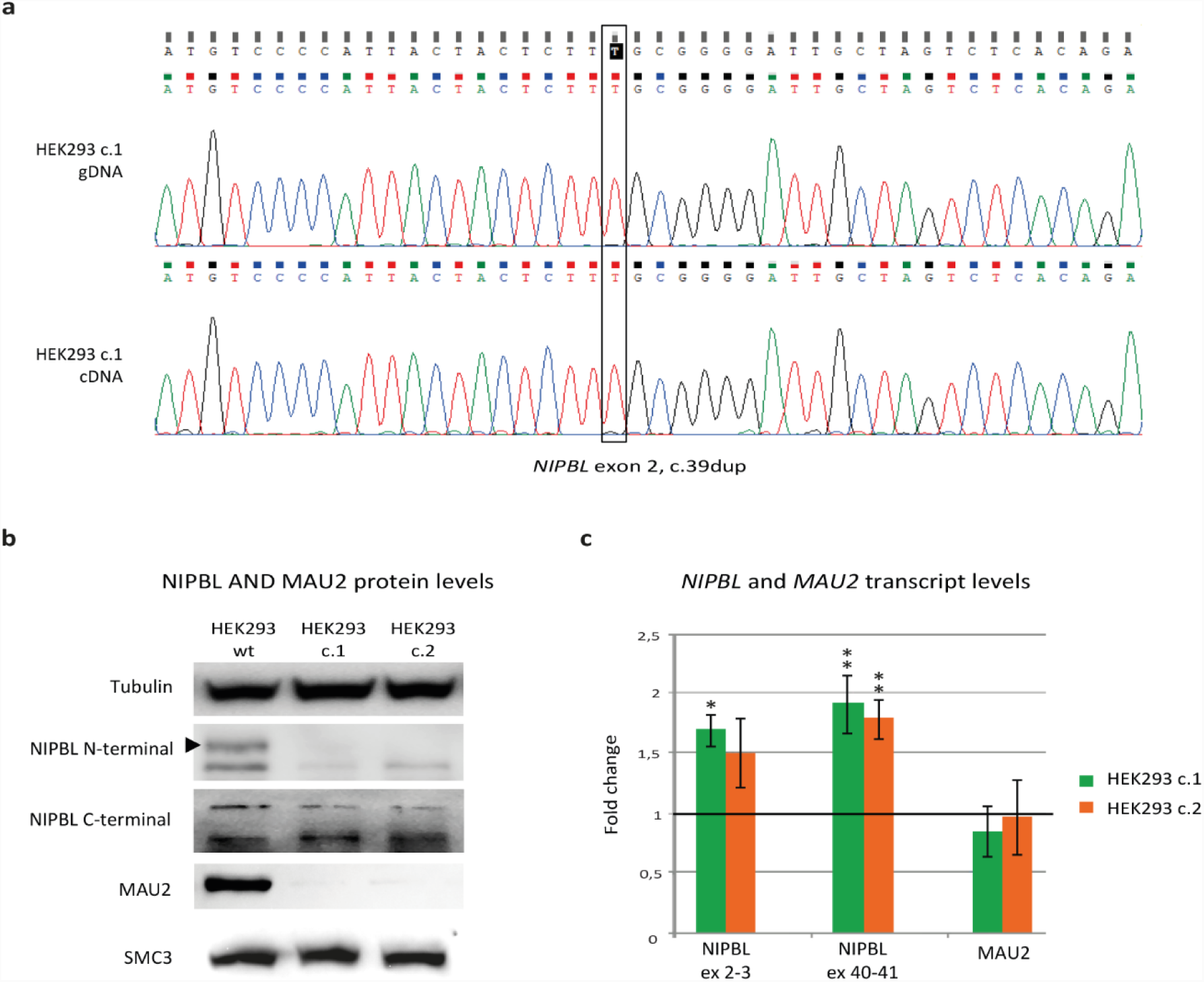
Genome edited cells express an N-terminal truncation of NIPBL and lack MAU2 a) Electropherograms of genomic DNA and cDNA of clone 1 (c.1) of the HEK293 edited cells, showing a homozygous duplication of T in exon 2 of *NIPBL*. b) Immunoblot analysis of NIPBLΔN cells: detection of NIPBL was carried out with two different antibodies, mapping to the N-terminus and C-terminus of NIPBL, as indicated. A reduced NIPBL signal could be detected only with the C-terminal antibody, indicating that an N-terminally truncated form of NIPBL is present in the CRISPR/Cas9 cells (NIPBLΔN cells). Additionally, the immunoblot analysis showed loss of MAU2, whereas there are no differences in the total amount of the cohesin subunit SMC3 between wild type and NIPBLΔN cells. c) Analysis of *NIPBL* and *MAU2* transcript levels: expression data were normalized to *GAPDH* and relative mRNA levels were determined using the ΔΔCt method. qPCR results indicate that *MAU2* expression is mainly unchanged in NIPBLΔN cells in comparison to its expression in wild type cells (set as 1 and represented as a horizontal black line) (Bilateral unpaired T-test; c1 p=0.09, c2 p=0.85). For *NIPBL* analysis, two different primer pairs were used, one of which was mapping between exons 2 and 3 and the other mapping between exons 40 and 41. Both assays show an upregulation of the *NIPBL* transcript in edited cells in comparison to wild type cells (Bilateral unpaired T-test; exon 2-3: c1 *p=0.03, c2 p=0.06; exon 40-41: c1 **p=0.006, c2 **p=0.001).

Immunoblot experiments were performed on the genome-edited cells in order to examine the expression of the kollerin complex subunits NIPBL and MAU2 and of the cohesin subunit SMC3 (Figure 1b). NIPBL expression was assessed with two different antibodies, one raised against its N-terminus and one recognizing its C-terminus. Notably, mutant *NIPBL* transcripts partially undergo nonsense-mediated mRNA decay in cell lines of patients with heterozygous truncating variants in *NIPBL*; consequently, the corresponding truncated protein is unlikely to be detected^32,33^. In both HEK293 clones carrying the homozygous out-of-frame duplication in *NIPBL*, the antibody raised against the N-terminus of NIPBL could not detect any protein signal. However, we were able to detect NIPBL in the edited clones using the C-terminal antibody, suggesting that only the N-terminus of the protein is missing. These results indicate that an N-terminally truncated form of NIPBL (NIPBLΔN) is expressed in the CRISPR/Cas9 clones (thereafter named NIPBLΔN cells) possibly as a result of the employment of an alternative Translation Initiation Site (aTIS). NIPBLΔN was found to be less abundant than full-length NIPBL in the parental cells, suggesting a lower efficiency of the alternative translation or a reduced stability of NIPBLΔN.

Importantly, no immunoblot signal could be detected for MAU2, suggesting an almost total loss of MAU2 in NIPBLΔN cells. The analysis of the structural subunit SMC3 indicated instead that the total amount of cohesin is not altered in NIPBLΔN cells.

### *NIPBL* mRNA level is upregulated in NIPBLΔN cells

The identification of a NIPBLΔN protein in these genome-edited cells suggested that the mutant transcript escapes nonsense-mediated mRNA decay. To test this hypothesis, we first performed Sanger sequencing of the cDNA obtained from NIPBLΔN cells. By this, we were able to confirm the expression of the mutant allele in a homozygous state (Figure 1a).

To expand upon this observation, we measured *NIPBL* transcript levels in control and in NIPBLΔN cells by Real-Time PCR. For this analysis, we designed two different primer pairs that cover boundaries between exons 2 and 3 and between exons 40 and 41, respectively. Both assays revealed a significant upregulation of *NIPBL* expression in NIPBLΔN cells compared to parental cells. Specifically, expression levels of the 5’ of *NIPBL* mutant transcript (*NIPBL c.39dup*) were respectively 68% and 50% higher for clones 1 and 2 in comparison to those observed in wild type cells. Similarly, expression levels of the 3’ of *NIPBL c.39dup* were 90% and 77% higher than in parental cells for clones 1 and 2, respectively (Figure 1c).

Altogether, these results corroborate the hypothesis that the *NIPBL c.39dup* transcripts do not undergo nonsense-mediated mRNA decay in NIPBLΔN cells and, instead, suggest the existence of a feedback mechanism between NIPBL protein and transcript to fine-tune its expression in order to ensure proper execution of its functions. A similar compensation mechanism is also observed in cell lines of patients with mutations in *NIPBL*, where the wild type allele is frequently upregulated, presumably in order to compensate for the downregulation of the mutant transcript induced by nonsense-mediated mRNA decay^28,33^.

Subsequent analysis of *MAU2* revealed no significant differences in the total amount of *MAU2* transcript in NIPBLΔN cells in comparison to wild type cells (Figure 1c). We therefore speculated that *MAU2* is correctly transcribed and translated and then degraded because of its inability to interact with NIPBL. To evaluate the stability of MAU2, wild type and NIPBLΔN cells were treated with the proteasome inhibitor MG132, the lysosomal inhibitor NH4Cl or the autophagy inhibitor Bafilomycin. Inhibition of these protein degradation pathways could not restore the expression of MAU2 in NIPBLΔN cells, indicating that none of them is involved in MAU2 degradation (Supplementary Figure 1).

Altogether, these results indicate that the transcripts of the kollerin subunits are correctly expressed and that the full-length *NIPBLc.39dup* transcript is used as a template for the production of NIPBLΔN, which is able to ensure, at least partly, the NIPBL functions that are essential for cell viability and proliferation.

### Identification of alternative translation initiation sites

To address whether an alternative start site allows initiation of NIPBL translation in NIPBLΔN cells, we established an In Vitro Transcription-coupled Translation (IVTT) assay.

Specifically, a fragment of *NIPBL* cDNA comprising the first 651 bp was cloned into a pcDNA3.1B vector for IVTT using a reverse primer adding a 3xFLAG-tag 3’ to the coding sequence. Both wild type and mutant *NIPBL* cDNAs were cloned into this vector using the same cloning primers. Of importance here, the 3xFLAG-tag sequence carried by the reverse primer was designed in frame with the wild type sequence and its canonical ATG but not with the mutant sequence harbouring the duplication of T (Figure 2a). The resulting plasmids served as templates in IVTT reactions. Immunoblot experiments with an anti-FLAG antibody were then performed in order to detect the proteins of interest. As shown in Figure 2c, a positive FLAG signal was detected in the IVTT product of the wild type as well as of the mutant plasmid, indicating that the Reticulocyte Lysate System of the IVTT reaction is indeed using an alternative translation start site to initiate protein synthesis in the presence of the out-of-frame duplication. Notably, the wild type and mutant IVTT proteins displayed only a very small difference in size (less than 2 kDa), hence indicating that the alternative translation start site is located in close proximity to the canonical ATG. In this context, the employment of the first ATG after the duplication of T would represent the most plausible alternative. However, the first ATG is located 99 amino acids downstream of the duplication. Such a distance would result in a substantial difference in size between wild type and mutant IVTT products. Since no such size difference was observed between the two proteins, we concluded that the potential alternative start site had to be located upstream of the mutation and hypothesised that it results from a shift of the canonical reading frame. In support of this hypothesis, we identified two new alternative Translation Initiation Sites (aTIS) after shifting the reading frame of one base pair (aTIS1 and aTIS2, Figure 2b). To determine which of these two putative aTISs was actually used, we subsequently inserted mutations in each aTIS (ATG>ACG) and performed IVTT using the resulting mutated plasmids (Figure 2c). A positive FLAG signal was detected after mutagenesis of aTIS1, whereas the signal was dramatically reduced in the IVTT product with a mutated aTIS2. These results indicate that aTIS2 is responsible for the initiation of translation of NIPBL in the presence of the out-of-frame duplication of T. The weak FLAG signal detected upon mutagenesis of aTIS2 suggests an inefficient usage of aTIS1. In support of this hypothesis, the FLAG signal was lost upon mutagenesis of both aTIS.

**Figure 2.**
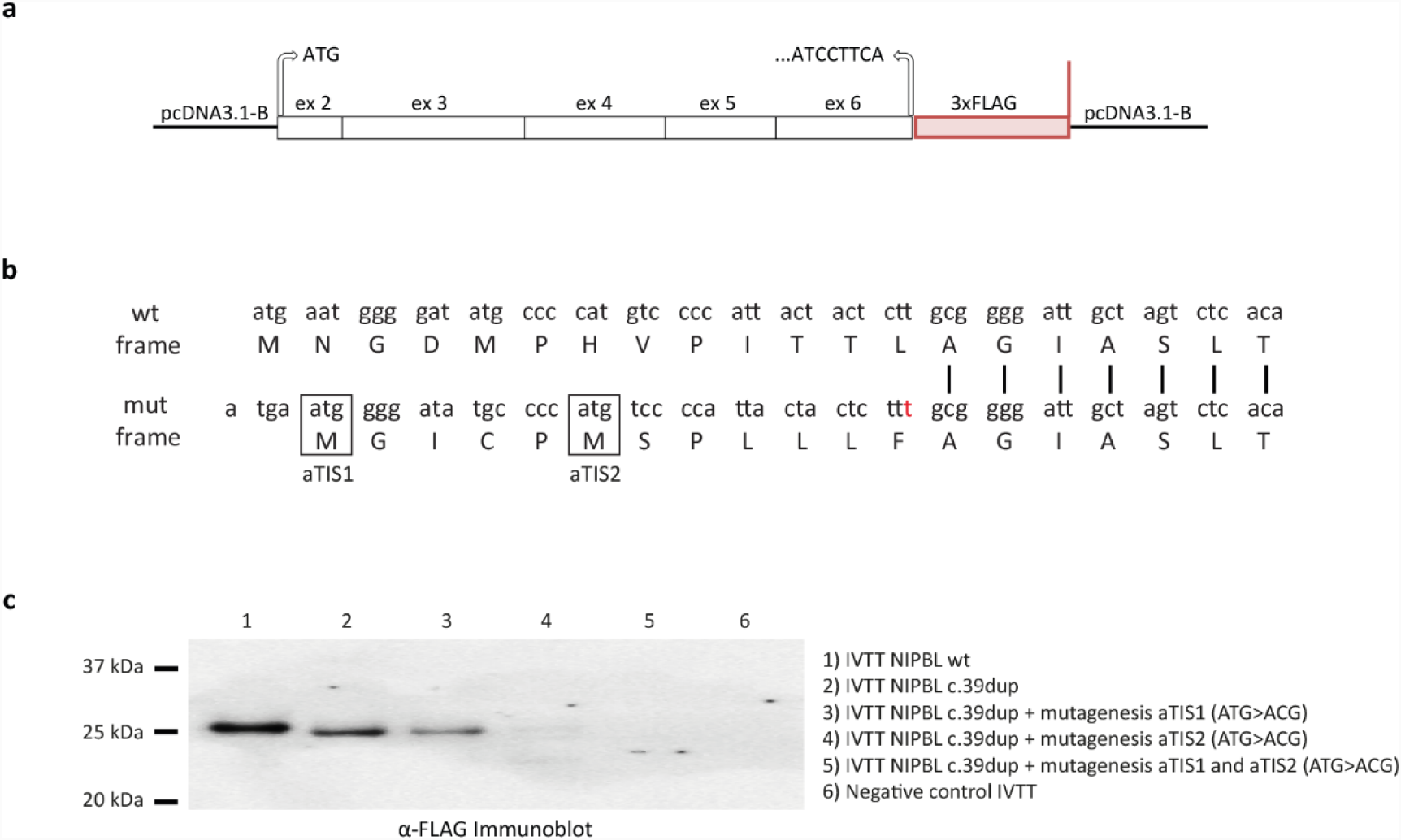
Identification of an alternative Translation Initiation Site a) To prove that an alternative translation start site is used in NIPBLΔN cells, we cloned the first 651 bp of *NIPBL* into a vector suitable for IVTT (pcDNA3.1B). Both wild type and mutant sequence of *NIPBL* were cloned using a reverse primer containing a 3xFLAG-tag that was designed in frame with the wild type sequence, but not with the mutant sequence carrying the duplication of T. b) An N-terminally truncated form of NIPBL (NIPBLΔN) is produced in the CRISPR/Cas9 cells in the presence of the c.39dup mutation. A very small difference in size was observed between the wild type and the mutant proteins by Immunoblot analysis, indicating that the alternative Translation Initiation Site (aTIS) should be in close proximity to the canonical ATG. The analysis of the sequence surrounding the duplication of T (red) suggested that two new ATGs appear after shifting the reading frame of one base pair (aTIS1 and aTIS2). The employment of one of these new ATGs to initiate translation would restore the correct frame of the protein after the duplication of T, producing an NIPBL isoform characterized by a different N-terminus and a correct C-terminus. Plasmids carrying mutations of the new putative aTIS, in addition to the duplication of T, were also generated. c) All plasmids were used as templates in IVTT reactions and the IVTT products were subsequently analysed by Immunoblot with a FLAG antibody. By this, we could detect a FLAG signal from the IVTT products of the wild type plasmid (1) as well as of the mutant one (2), confirming that an alternative translation start site is used in the presence of the duplication of T. The FLAG signal was still detectable after mutagenesis of aTIS1 (3), but was dramatically downregulated after mutagenesis of aTIS2 (4), indicating that, in the presence of the duplication of T, aTIS2 is used to start the translation of NIPBLΔN. The weak FLAG signal observed in the Immunoblot of the IVTT product 4 is probably associated with inefficient usage of aTIS1. Accordingly, the FLAG signal was lost after mutating both aTIS (5).

The employment of aTIS2 to initiate translation of NIPBLΔN in the edited cells restores the correct frame of the protein after the duplication of T, producing an isoform of NIPBL characterized by a different N-terminus and a correct C-terminus. Precisely, only the first 12 amino acids differentiate the wild type NIPBL from the new isoform. However, this alternative isoform is predicted to be unable to interact with MAU2, since the first 38 amino acids of NIPBL are responsible for this interaction^9^. Consistently, molecular characterization of NIPBLΔN cells showed MAU2 depletion, most likely due to its inability to interact with NIPBL.

### Cohesin-binding to chromatin is not affected in NIPBLΔN cells

To measure the effects of MAU2 deficiency on the abundance and dynamics of NIPBLΔN and cohesin on DNA, we first fractionated wild type and NIPBLΔN cells (clones c.1 and c.2) into total, soluble and chromatin-bound fractions (Figure 3a). A modest reduction of NIPBL and the loss of MAU2 were observed in all fractions of both NIPBLΔN clones. No difference in the total amount of chromatin-bound cohesin was observed between the two cell lines, indicating that cohesin complexes are still able to bind chromatin in NIPBLΔN cells despite the absence of MAU2 and in the presence of an N-terminally truncated version of NIPBL.

**Figure 3.**
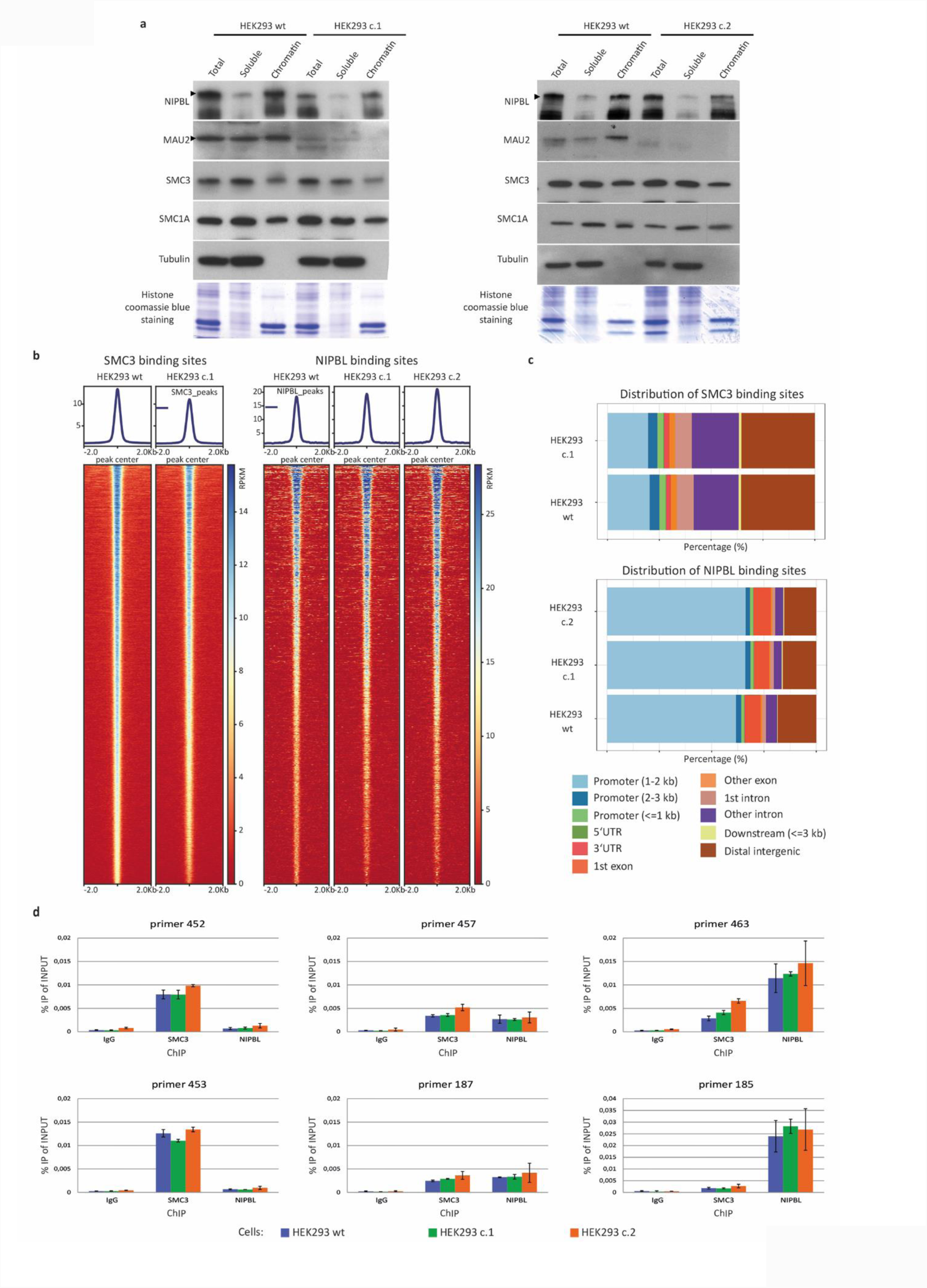
Cohesin-binding to chromatin is not affected in NIPBLΔN cells a) Fractionation of wild type and NIPBLΔN cells (c.1 and c.2) in soluble and chromatin-bound proteins shows a small reduction of NIPBL and loss of MAU2 in the soluble pool but also on chromatin in NIPBLΔN cells. No differences in the amounts of soluble or chromatin-bound cohesin are observed. b) The heatmaps depict normalised ChIP coverage observed in the SMC3 and NIPBL ChIP-seq experiments performed with formaldehyde crosslinking for wild type and NIPBLΔN cells. The heatmaps are centred on the peaks observed in wild type cells and the colour intensity relates to normalised counts (RPKM). c) Distribution of the peaks observed in the SMC3 and NIPBL ChIP-seq experiments over different genomic features (eg. promoters, introns, exons) for wild type and NIPBLΔN cells. No striking differences are visible between wild type and NIPBLΔN cells. d) Several strong binding sites for cohesin (primers 452 and 453) and NIPBL (primers 463 and 185) as well as two sites weakly bound by cohesin and NIPBL (primers 457 and 187) were analysed by ChIP-qPCR in wild type cells and NIPBLΔN cells. This analysis indicates that the amount of chromatin-bound cohesin and chromatin-bound NIPBL are mainly unchanged between wild type and NIPBLΔN cells at these sites. More sites can be found in Supplementary figure 3.

Next, we performed chromatin immunoprecipitation followed by sequencing (ChIP-seq) for NIPBL and the cohesin subunit SMC3. We called 23,194 SMC3 peaks in wild type cells and 21,984 peaks in NIPBLΔN cells (Figure 3b and Supplementary Figure 2a) and the heatmaps comparing the peaks signals in wild type cells with the signal intensity in NIPBLΔN cells showed a slightly reduced signal in the NIPBLΔN cells, an effect that is more pronounced at weaker peaks. The distribution of the peaks over different genomic features, however, is unchanged (Figure 3c). For NIPBL we called 2,681 peaks for wild type cells and 2,505 and 2,695 peaks for NIPBLΔN clones 1 and 2, respectively (Figure 3 b-c, Supplementary Figure 2b). The identified peaks do not show a change in feature distribution. Upon application of a ChIP-seq protocol involving protein-protein-crosslinking in addition to formaldehyde crosslinking to pick up more efficiently weaker NIPBL binding sites, we were able to call 14,198 peaks in wild type cells and 9,124 peaks in NIPBLΔN cells (Supplementary figure 2b). Heatmaps comparing the peak signals in wild type cells with NIPBLΔN cells (Supplementary Figure 2c) show that on all wild type peaks we can still obtain signal in NIPBLΔN cells, indicating that a large number of the peaks in NIPBLΔN is still present but just dropped under the peak cutoff. This is also supported by the notion that the overall distribution of the peaks over different genomic features is not altered.

The results obtained by ChIP-seq were subsequently confirmed by ChIP-qPCR experiments, where SMC3 and NIPBL binding was assessed at different genomic loci. These analyses confirmed that cohesin- and NIPBL-binding sites are largely unchanged in NIPBLΔN cells (Figure 3d; Supplementary Figure 3).

Altogether, these experiments indicate that NIPBLΔN is able to bind chromatin and to mediate normal cohesin loading onto DNA despite the absence of MAU2. However, although the position of binding sites along the genome is not altered, a lower occupancy was detected, consistent with the observed reduction of the chromatin-bound NIPBL fraction.

### NIPBLΔN cells do not display cohesion defects

NIPBL is well known for its role as cohesin loader. The canonical function of the cohesin complex is the maintenance of sister chromatid cohesion during both mitosis and meiosis.

In CRISPR/Cas9 cells, NIPBLΔN is downregulated in comparison to parental cells, but cohesin appears to be efficiently loaded onto the DNA. To address whether the chromatin-bound cohesin is able to fulfil its cohesive function, we analysed NIPBLΔN and control cells by spreading and Giemsa staining of mitotic chromosome after two hours treatment with the spindle poison nocodazole. In both NIPBLΔN clones and control cells, a normal cohesion was observed in more than 85% of the analysed mitotic cells, indicating the absence of evident cohesion defects in NIPBLΔN cells (Figure 4a, b). In addition, determination of mitotic indices revealed no significant differences between NIPBLΔN and parental cells (Figure 4c).

**Figure 4:**
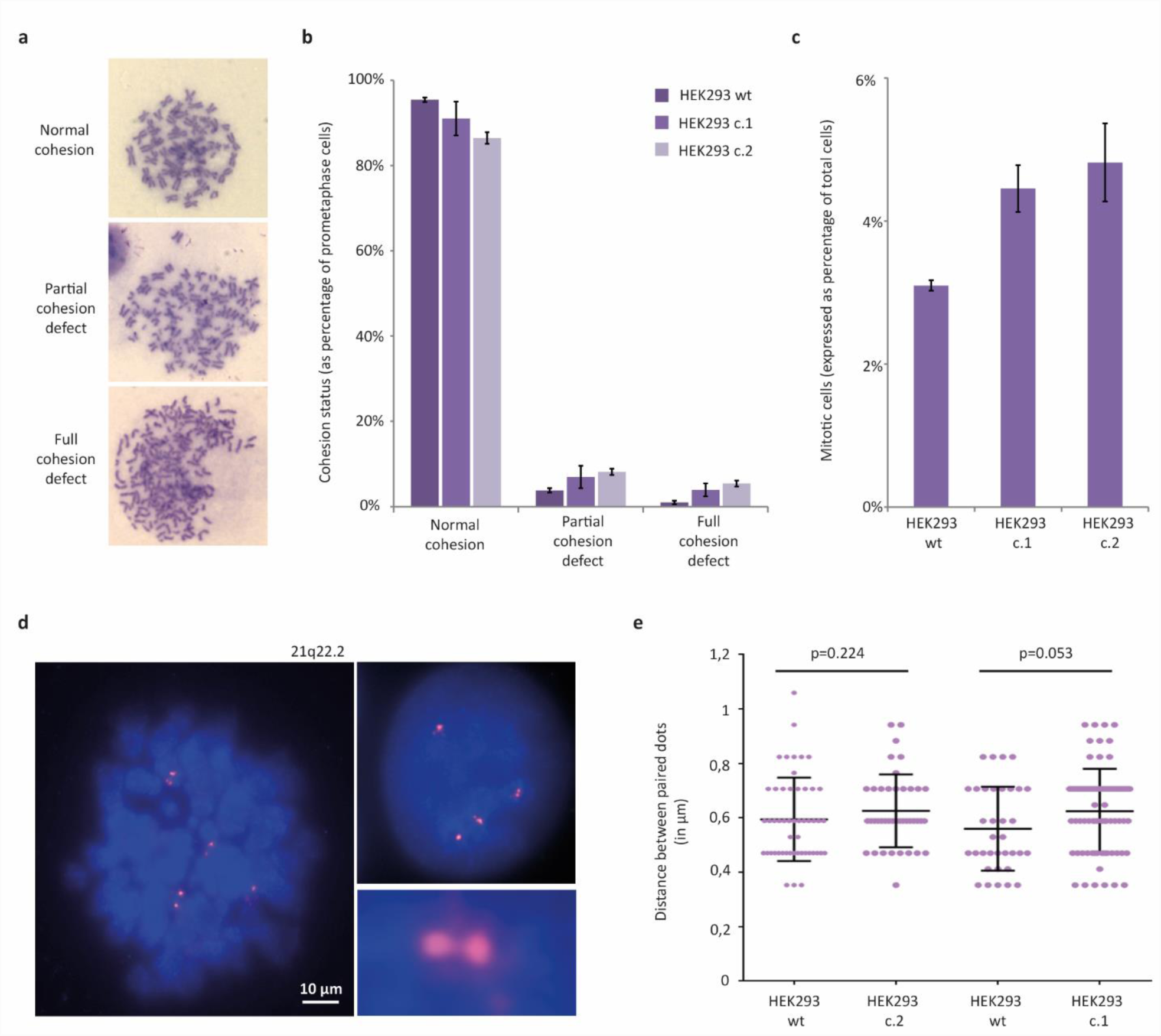
NIPBLΔN cells do not display cohesion defects a) Representative images of chromosome spreads. Cohesion was considered normal if chromosomes contained a primary constriction, i.e., sister chromatids were tightly connected at their centromeres (upper panel). Chromosomes that lacked a primary constriction and in which sister chromatids were abnormally spaced, but in which chromatids were still closely opposed to each other, were considered to have mild cohesion defects (middle panel). Complete separation of sister chromatids indicated loss of cohesion (bottom panel). b) Graphical representation of the frequency of the different cohesion phenotypes was determined and expressed as a percentage of total prometaphase cells. No increased frequency of cohesion defects was detected during mitosis in the presence of NIPBLΔN and in the absence of MAU2. c) Graphical representation of the mitotic index of wild type and NIPBLΔN cells, expressed as percentage of total cells. d) Representative image of interphase DNA FISH performed on NIPBLΔN cells. Bar represents 10 µm. e) Graphical representation of the quantification of the interphase DNA FISH, showing that the distance between paired FISH signals is not affected, confirming the absence of cohesion defects.

Further analysis of sister chromatid cohesion in these cells was performed by measuring the distance between paired FISH signals in interphase cells. We used a FISH probe that maps on chromosome 21q22.2, a locus that is tetrasomic in HEK293 cells, and therefore labels four pairs of sister chromatids in G2 cells (Figure 4d). In wild type cells, the average interchromatid distance was 0.57 μm. The presence of NIPBLΔN in both *NIPBL c.39dup* clones did not change the distance between FISH signals significantly (Figure 4e). Altogether, these results indicate that there are no sister chromatid cohesion defects during G2 phase and mitosis in the presence of NIPBLΔN and in the absence of MAU2.

### Whole genome sequencing identifies the first mutation in *MAU2*

Trio whole genome sequencing was performed on a group of 15 patients with a clinical diagnosis of CdLS who were previously found to be negative for the presence of mutations in the five known CdLS genes. This analysis led us to the identification of the first putative pathogenic variant in *MAU2*, detected in a patient with a characteristic CdLS phenotype (Figure 5a). The patient was born at 34 weeks of gestation after caesarean section from healthy and non-consanguineous parents. Intra-uterine growth retardation was detected at the 30^th^ week of pregnancy. At birth, Apgar scores were 5 and 8 at 1 and 5 minutes, respectively. Birth weight was 1330 g (−2.46 SD), birth length was 39.5 cm (−2.42 SD) and the occipital frontal circumference was 27 cm (−3.17 SD). He showed early feeding difficulties and developed a generalized spasticity from the very first weeks of life, which severely affected his mobility and soon required the use of a wheelchair. At the age of two years he was diagnosed with Cornelia de Lange syndrome. His facial features were typical for the syndrome and included microcephaly, brachycephaly, low anterior and posterior hairline, arched eyebrows, synophrys, long eyelashes, ptosis, flat nasal bridge, long and smooth philtrum, thin lips with downturned corners of the mouth and highly arched palate. In addition, he presented with hypertelorism, myopia, low-set and posteriorly rotated ears, small feet, clinodactyly of the fifth finger, hirsutism and cryptorchidism. He developed very early gastroesophageal reflux disease. Magnetic Resonance Imaging revealed the existence of a thin corpus callosum, a mild ventriculomegaly and periventricular cysts. Last evaluation was performed at the age of 10 years. At this age, weight was 18.8 kg (−4.5 SD), height was 121 cm (−4.96 SD) and the occipital frontal circumference was 47 cm (−4.96 SD). He presented with severe intellectual disability and delayed speech and motor development. He pronounced his first words at the age of two years but had no further verbal development. Currently, he is still not able to sit unassisted or to walk.

**Figure 5.**
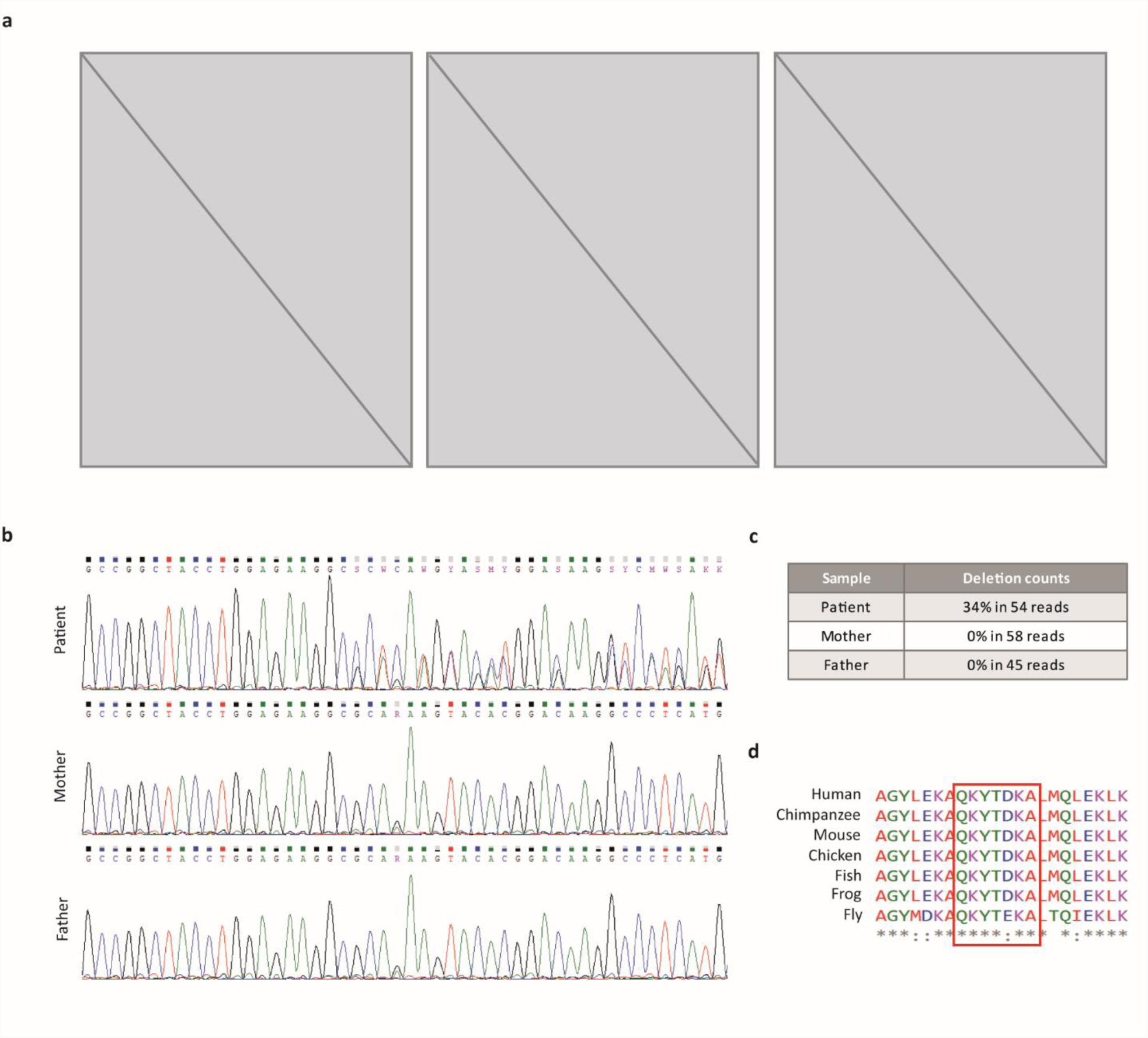
Identification of a *MAU2* mutation in a patient with CdLS a) Phenotypical appearance of the CdLS patient carrying the in-frame deletion in *MAU2* (c.927_947del; p.(Gln310_Ala316del)). The first two pictures were taken at the age of two years, while the picture on the right was taken at the age of 10 years. Please note the typical CdLS facial features and evident spasticity (<u>Pictures were removed because the BioRxiv policy does not allow the inclusion of photographs or any other identifying information of people</u>). b) Sanger sequencing of the blood DNA of the patient and of its parents, illustrating the *de novo* origin of the mutation. c) The deletion counts of the genome sequencing run indicates the percentage of mutant allele in the blood DNA of the patient and of his parents and confirms the *de novo* origin of the mutation. d) Alignment of multiple orthologues of MAU2. A section of exon 9 of MAU2 was aligned in seven different species (*Homo sapiens, Pan troglodytes, Mus musculus, Gallus gallus, Esox lucius, Xenopus laevis* and *Drosophila melanogaster*). The amino acids residues affected by the deletion, highlighted in the red square, show a high level of conservation from humans to flies.

This patient was found to carry an in-frame deletion of 21 nucleotides in *MAU2*, resulting in the loss of seven amino acids: RefSeq NM_015329, c.927_947del, p.(Gln310_Ala316del). The variant was identified in 34% of the sequencing reads, and Sanger sequencing could confirm its *de novo* origin (Figure 5b, c). Alignment of the MAU2 protein sequence across seven species highlighted that the deletion affects a region that is highly conserved in both vertebrate and non-vertebrate animal species (Figure 5d). To date, *de novo* missense and nonsense mutations in *MAU2* have been found to be significantly underrepresented in exome sequences^10^. The Exome Aggregation Consortium (ExAC) categorizes *MAU2* among the haploinsufficient genes^34^. The pLI (probability of intolerance) score of *MAU2* calculated by the database is in fact equal to 1, which reveals that the gene is extremely intolerant to loss-of-function variations. Similarly, the Z score of *MAU2* is equal to 4.92. This score estimates the ability of a gene to tolerate missense substitutions. A positive Z score, such as the one associated to *MAU2*, indicates that the gene has fewer variants than expected, implying increased constraint towards variations. Taken together, these data strongly suggest that the mutation identified in the patient of our cohort represents the first pathogenic variant in *MAU2*.

### The deletion in MAU2 impairs the heterodimerization with NIPBL

To investigate how the identified deletion could affect MAU2 protein structure and its ability to bind NIPBL, we performed an *in silico* analysis of the co-crystal structure of the *S. cerevisiae* cohesin loader complex Scc2/NIPBL-Scc4/MAU2 (4XDN)^10^.

Based on a previous comparison of the MAU2/Scc4 protein sequence from yeast to human^5^ we mapped the deleted region on the Scc2/NIPBL-Scc4/MAU2 crystal structure from *S. cerevisiae* (Figure 6a). The deletion affects a helix that is part of the TPR array, a domain that envelops the N-terminus of NIPBL. The deletion removes part of a helix that contacts directly the Scc2/NIPBL N-terminus, likely leading to a distortion of the MAU2-NIPBL interaction (Figure 6b).

**Figure 6.**
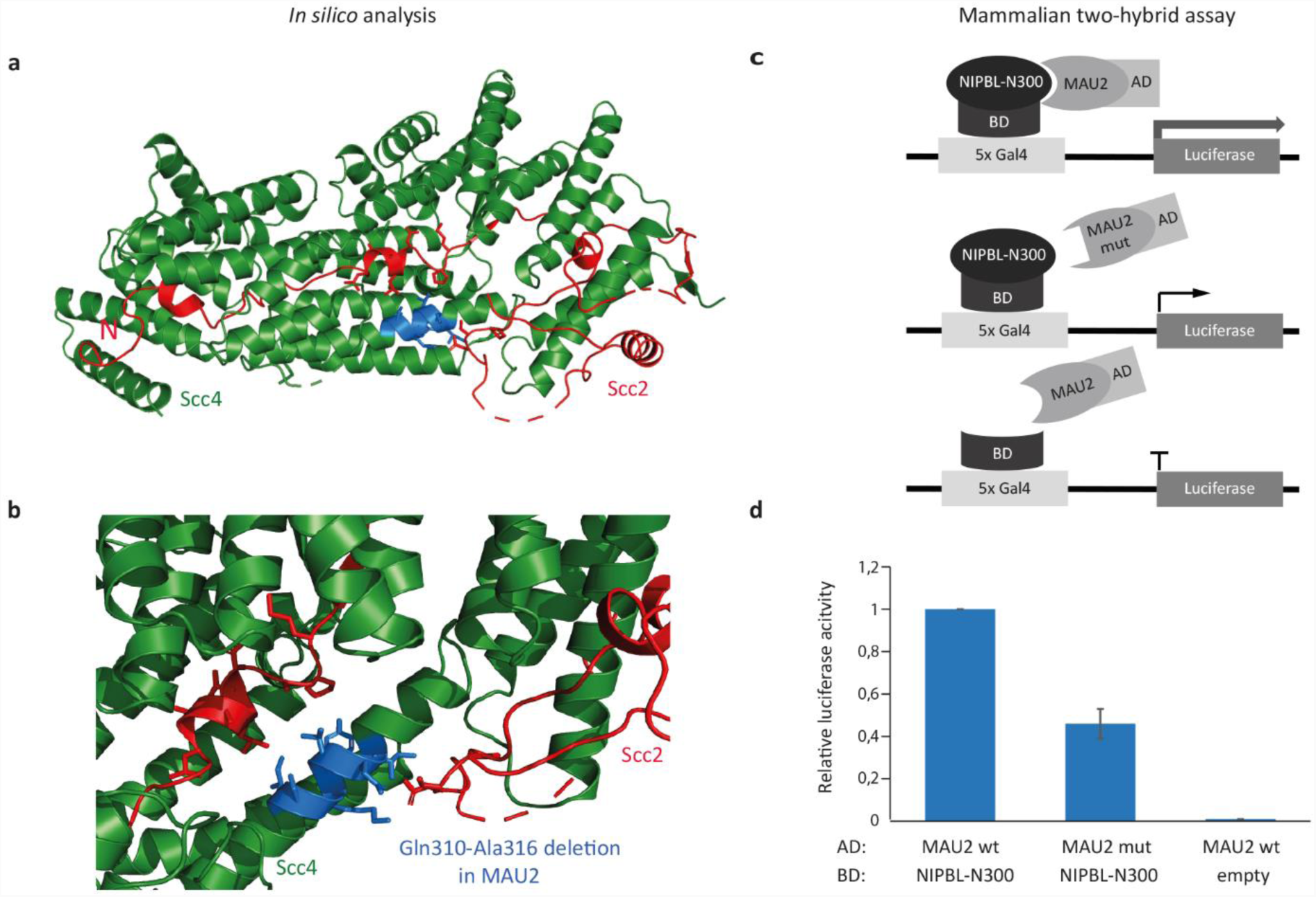
The in-frame deletion p.(Gln310_Ala316del) in MAU2 impairs its interaction with NIPBL. a-b) *In silico* analysis of the co-crystal structure of the *S. cerevisiae* cohesin loader (4XDN)^10^. The N-terminus of Scc2/NIPBL (depicted in red) is enveloped by the TPR repeats of Scc4/MAU2 (depicted in green). The helix fragment in the *S. cerevisiae* structure corresponding to the deletion Gln310-Ala316 of the human protein is depicted in light blue. The deletion is likely to distort the interactions between Scc4/MAU2 and Scc2/NIPBL (a). Besides the overall distortion of the structure by the shortening of the helix, the deleted residues are likely interacting with residues close to the N-terminus of Scc2/NIPBL but also loops further into the structure (b). c-d) Mammalian two-hybrid assay was performed using the full length MAU2 wild type (wt) or mutant (mut) conjugated to the activation domain (AD) of the mouse NF-κB and the first 300 amino acids of NIPBL (NIPBL-N300) conjugated to the binding domain (BD) of the GAL4 gene. The BD domain alone was instead used as negative control. The results obtained indicate that the *MAU2* in-frame deletion p.(Gln310_Ala316del) causes a reduction of the interaction between NIPBL and MAU2 of approximately 50%.

To test this prediction, we introduced the mutation into plasmids to perform quantitative mammalian two-hybrid interaction assays in HEK293 cells. Constructs encoding the N-terminal 300 amino acids of NIPBL coupled to the DNA binding domain of Gal4 were tested for interaction with the wild type or mutant full-length MAU2, bound to the activation domain of NF-κB (Figure 6c). This assay demonstrated that the seven amino acid deletion in MAU2 causes a reduction of the heterodimerization activity to 46% (Figure 6d). The deleterious effect of MAU2 deletion on its interaction with NIPBL was further confirmed with the yeast two-hybrid assay. Indeed, yeast transformed with mutant MAU2 showed a slower growth on selective medium, indicative of a weaker interaction with NIPBL in comparison to wild type MAU2 (Supplementary Figure 4). Taken together, these *in vitro* interaction assays support the *in silico* predictions and demonstrate that the MAU2 deletion impairs the interaction of the two subunits of the kollerin complex.

## DISCUSSION

*NIPBL* is an ubiquitously expressed gene that is essential for human development^16^. The resulting protein interacts with MAU2 in a complex named kollerin, which is involved in transcriptional regulation and cohesin loading^4,6^. Mutations in *NIPBL* are the primary genetic cause of CdLS, whereas alterations of its binding partner *MAU2* have never been reported so far^35^. CdLS is characterized by an extensive clinical variability and patients present with a wide range of phenotypes^17^. Heterozygous out-of-frame deletions and insertions or nonsense variants in *NIPBL* are mainly associated with severe phenotypes and with the presence of limb malformations^20^. However, truncating mutations affecting exons 2 to 9 of *NIPBL* are associated with a lower frequency of limb reductions or malformations and often result in phenotypes that are milder than expected in comparison to truncating mutations affecting exons 11-47^18,30,31^. Our genome editing experiments demonstrated that cells with early truncations in *NIPBL* adopt a protective mechanism based on alternative translation initiation in order to minimise the otherwise deleterious effects of mutations, thus offering an explanation for the milder phenotypes of these patients. The choice of the new translation initiation site depends on the nucleotide sequence surrounding the variant of interest, thereby rendering this protective mechanism mutation-specific. By this, cells ensure the synthesis of the essential C-terminus of NIPBL at the expenses of its N-terminus, which appears to be dispensable for cell survival. Congruent with this hypothesis, Cap Analysis of Gene Expression (CAGE) data suggest the existence of a possible alternative Transcription Start Site (TSS) in exon 10 of *NIPBL* (http://fantom3.gsc.riken.jp/). In addition, a recent paper demonstrated that cells could tolerate disruptive mutations up to exon 10, indicating that the region between exons 11 and 47 alone is able to accomplish the main tasks of NIPBL and is therefore essential for cell survival^36^. Herein, we described viable cells expressing an N-terminally truncated form of NIPBL (NIPBLΔN) that lacks the first 12 amino acids. Since the first 38 amino acids of NIPBL are important for the interaction with MAU2^9^, NIPBLΔN is unable to interact with MAU2 in the genome-edited cells, leading to MAU2 depletion. Molecular analyses of these cells indicated that NIPBLΔN is still able to bind chromatin and to mediate cohesin loading onto DNA despite the absence of MAU2. This notion is in line with recent findings stating that a mutant NIPBL construct that is incapable to interact with MAU2 is still able to stimulate the ATPase activity of the cohesin complex and to mediate DNA repair^37,38^. Similarly, Murayama and Uhlmann formerly reported that *in vitro* NIPBL alone displayed DNA binding properties indistinguishable from that of the kollerin complex and was able to mediate cohesin loading onto chromatin^2^.

In our cells, cohesin and NIPBL binding sites are largely unchanged. Chromatin-bound cohesin also proved to be able to accomplish the main function of the complex, as cohesion defects were not detected in NIPBLΔN cells. Correspondingly, cell line of patients with a clinical diagnosis of CdLS do not display cohesion defects^39^. However, previous data suggested that a particular patch on the MAU2 surface is important for cohesin loading to the centromere in yeast^10^. Our findings therefore suggest that the interaction between MAU2 and cohesin is not necessary for sister chromatid cohesion in mammalian cells.

Altogether, these data provide further characterisation of the interaction between the two subunits of the kollerin complex. Downregulation of NIPBL by RNA interference was previously found to reduces MAU2 levels^5^. In line with this notion, MAU2 levels are strongly downregulated in NIPBLΔN cells as a consequence of the loss of the MAU2-interacting N-terminus domain of NIPBL. Similarly, in the presence of a full-length NIPBL, MAU2 is required for the correct folding of its N-terminus and for its stability^7^. Accordingly, the NIPBL missense substitutions p.(Gly15Arg) and p.(Pro29Gln), that alter the NIPBL-MAU2 interaction domain, induce a substantial reduction of the heterodimerization activity of the two proteins, thus leading to NIPBL haploinsufficiency and to a CdLS phenotype^9^.

In this paper, we describe the first mutation in *MAU2* that results in a CdLS phenotype. Similarly to the aforementioned missense mutations in *NIPBL*, the p.(Gln310_Ala316del) variant identified in the patient of our cohort leads to a distortion of the protein structure that impairs the interaction between the two subunits of the kollerin complex, hence affecting their stability. For this reason, we propose that this deletion may exert its deleterious effects in a way that is comparable to mutations that directly affect the coding sequence of *NIPBL* and that leads to *NIPBL* haploinsufficiency. In support of this hypothesis, the phenotype of our patient resembles the phenotype of patients with mutations in *NIPBL*. In particular, this patient presents with typical CdLS facial dysmorphism, microcephaly, severe intellectual disability and with an extremely delayed motor and verbal development, all features pointing to a classical manifestation of the syndrome. A particular resemblance stands out when comparing the phenotype of our patient with the patient carrying the p.(Gly15Arg) substitution in *NIPBL*^9^. Both patients present with a low anterior hairline, synophrys, long eyelashes, ptosis, a flat nasal bridge, long and featureless philtrum and thin lips. Notably, both individuals also show macrodontia of central incisors, a feature per se not typical for CdLS. This resemblance supports the hypothesis that the deletion in *MAU2* and the missense mutation in *NIPBL* might be associated with similar downstream molecular consequences.

Altogether, our data confirm NIPBL haploinsufficiency as the major pathogenic mechanism of CdLS but also give new insights into the molecular mechanisms responsible for this neurodevelopmental disorder. Specifically, our functional investigations point to a CdLS relevant function of NIPBL that is not related to cohesin loading and reveal the existence of a new pathogenic mechanism resulting in NIPBL instability upon functional alteration of its binding partner MAU2.

Our data also strongly indicate the existence of a protective mechanism preventing a total loss of NIPBL gene product by the use of alternative start codons in transcripts with early truncating mutations. To our knowledge, this is the first report of such a protective mechanism, and it might as well be possible that a similar alternative translation start-based rescue is employed in the presence of loss-of-function mutations in other dosage-sensitive genes responsible for neurodevelopmental disorders, hence opening perspectives for translational approaches.

## Acknowledgments

We thank Jaulin’s team for access to the microscope, Koichi Tanaka and Kim Nasmyth for sharing the NIPBL antibody and Jan-Michael Peters for sharing the MAU2 antibody. This work was funded by the German Federal Ministry of Education and Research (BMBF) (CHROMATIN-Net: to F.J.K.), by E-Rare-2 TARGET-CdLS (to F.J.K, K.S.W. and E.W.), by the NWO Building blocks of life program project “Genometrack” (to K.S.W) and by the Medical Faculty of the University of Lübeck (J09-2017 to I.P.).

## MATERIALS AND METHODS CRISPR/Cas9

HEK293 cells were cultured in DMEM supplemented with 10% FBS and 1% Antibiotics (Thermo Fisher Scientific, Darmstadt, Germany) at 37°C with 4% CO2. The genome editing was performed as described by Ran and colleagues in 2013^40^. In detail, two different 20 nucleotides guide sequences followed by a 5’-NGG PAM and complementary to exon 2 of *NIPBL* (Biomers, Ulm, Germany, primers available upon request) were inserted into the pSpCas9(BB)-2A-Puro (PX459) V2.0 vector (#62988, Addgene, Cambridge, MA, USA). HEK293 cells were transfected with the resulting plasmids with the TaKaRa Xfect™ Transfection Reagent (Clontech-Takara, Saint-Germain-en-Laye, France), following the manufacturer’s instructions. 24 hours post-transfection, selection of positively transfected clones was performed for 48 hours with DMEM medium supplemented with 10% FBS, 1% Antibiotics and puromycin at a final concentration of 8 µg/ml (Thermo Fisher Scientific). After selection, single clones were picked and expanded. DNA of clones was isolated with the innuPREP DNA Micro Kit (Analytik Jena, Jena, Germany) according to the manufacturer’s protocol. Sanger sequencing of the region of interest was performed with the Big Dye terminator v3.1 Sequencing Kit and run on the ABI PRISM 3130xl Genetic Analyzer (Thermo Fisher Scientific). Electropherograms were analyzed with the SeqManII software version 5.06 (DNASTAR).

### Protein isolation, quantification, and Immunoblots

Cells were lysed using RIPA buffer ph 7.6 (50 mM HEPES, 1 mM EDTA, 1% NP-40, 0.5M LiCl) and a protease inhibitor cocktail (Roche, Mannheim, Germany). Protein concentration was determined through the BCA Protein Assay Kit (Pierce, Rockford, IL, USA) according to the manufacturer’s protocol with the Tristar^2^ Multimode Reader LB942 (Berthold Technologies, Bad Wildbad, Germany). For each sample, 20 µg of proteins were supplemented with Laemmli buffer (60 mMTris-Cl pH 6.8, 2% SDS, 10% glycerol, 5% β-mercaptoethanol, 0.01% bromophenol blue) and denatured at 95°C for 5 minutes. Protein transfer was performed at 25 V for 30 minutes with the Trans-Blot Turbo Blotting system (Bio-Rad, Hercules, CA, USA). Non-specific binding was blocked by incubating the membranes in 4% skimmed milk in PBS for 1 hour at room temperature.

Membranes were incubated overnight at 4°C in agitation with the respective primary antibodies. The incubation with the secondary antibodies was performed in 4% skimmed milk in PBS for 90 minutes at room temperature. Bound antibodies were detected using the SuperSignal™ West Femto Maximum Sensitivity Substrate (Thermo Fisher Scientific). Blot images were acquired with the MF-ChemiBIS 2.0 (Bio-Imaging Systems, Neve Yamin, Israel).

### Antibodies

For Western blot detection all antibodies were diluted 1:1000 in 4% skimmed milk in PBS. The following antibodies were used for the analysis: rabbit-α-MAU2 (ab183033, Abcam, Cambridge, MA, USA); rabbit-α-MAU2 (gift from Jan-Michael Peters); rat-α-NIPBL (010702F01, Absea Biotechnology, Beijing, China); mouse-α-NIPBL (sc-374625, Santa Cruz Biotechnology, Dallas, TX, USA); rabbit-α-SMC1A (ab21583, Abcam); rabbit-α-RAD21 (#4321, Cell Signaling Technology, Danvers, MA, USA); mouse-α-tubulin (T5201, Sigma-Aldrich, St. Louis, MO, USA); mouse-α-GAL4 (DBD) (sc-510, Santa Cruz); rabbit-α-NFκB antibody (sc-372, Santa Cruz); mouse-α-GAL4 (AD) (630402, Clontech); goat-α-rabbit (31460, Thermo Fisher Scientific); goat-α-mouse (31430 Thermo Fischer Scientific); goat-α-rat (31470, Thermo Fisher Scientific). For ChIP: Rabbit-α-SMC3 antibodies were obtained by immunizing rabbits using the C-EMAKDFVEDDTTHG peptide and subsequently purified using the peptide epitope coupled to a SulfoLink coupling resin (Thermo Fisher Scientific). Rabbit anti-NIPBL antiserum raised against residues 2598–2825 of the *X. laevis* Scc2-1B (133M) (gift from Koichi Tanaka and Kim Nasmyth) was used.

### RNA isolation, cDNA synthesis and Real-Time PCR

RNA extraction was performed with the ReliaPrep™ RNA Cell Miniprep System (Promega, Mannheim, Germany) according to the manufacturer’s instructions. Subsequent treatment with DNase I (RNase-free, New England Biolabs, Frankfurt am Main, Germany) was carried out on all RNA samples in order to avoid genomic DNA contaminations.

The SuperScript™ III Reverse Transcriptase (Thermo Fisher Scientific) was used to retro-transcribe 2 µg of RNA with random hexamers. cDNA synthesis was performed in two independent experiments for each sample.

The expression level of the transcripts of interest was assessed by the use of the Real-Time PCR qPCRBIO Probe Mix Hi-ROX assay (PCR Biosystems, London, UK). The investigation was run on the 7300 Real-Time PCR system (Thermo Fisher Scientific). The following TaqMan gene expression assays were used for the analysis: Hs01062386_m1, Hs00209846_m1 and Hs01122291_m1 (Thermo Fisher Scientific). Based on efficiency experiments, the GAPDH gene was selected as endogenous normalizer and amplified with the TaqMan gene expression assay ID Hs02758991_g1 (Thermo Fisher Scientific). Relative gene expression was determined using the ΔΔCt method, as previously described^41^.

### Protein degradation pathways

Wild type and CRISPR/Cas9 HEK293 cells were treated with the proteasome inhibitor MG132 for 4-8 hours (10 μM; Sigma Aldrich), with the lysosome inhibitor ammonium chloride for 3-6 hours (NH4Cl, 20 mM; Merck, Darmstadt, Germany) or with the autophagy inhibitor Bafilomycin for 3-6 hours (100 nM, Sigma Aldrich).

### *In vitro* transcription-coupled translation

In Vitro Transcription-coupled Translation (IVTT) reactions were performed with the TnT® Quick Coupled Transcription/Translation System (Promega) starting from 500 ng of plasmid, following the manufacturer’s instructions.

### Subcellular fractionation

HEK293 cells were collected off the plates and washed twice with ice-cold PBS. The final cell pellet was resuspended in extraction buffer (20 mMTris pH 7.5, 100 mM sodium chloride, 5 mM magnesium chloride, 0.2 % NP-40, 10 % glycerol, 0.5 mM dithiothreitol) supplemented with tablets of protease inhibitors (Roche) and phosphate inhibitors (10 mM sodium fluoride, 20 mM β-glycerophosphate). Cells were lysed on ice by passage through a 26-gauge needle. Lysates were incubated for 10 minutes on ice and were then centrifuged (21,000 g, 15 minutes, 4°C) for collection of the soluble protein extract. The chromatin-containing pellet was washed six times with extraction buffer (8,000 g, 3 minutes, 4°C) and then directly resuspended in sample buffer.

### Chip-sequencing

Chromatin immunoprecipitation for SMC3 was performed as previously described^42^. In brief, cells at 70–80% confluence were crosslinked with 1% formaldehyde for 10 minutes and quenched with 125 mM glycine. After washing with PBS, cells were resuspended in lysis buffer (50 mM Tris-HCl pH 8.0, 1% SDS, 10 mM EDTA, 1 mM PMSF and Complete Protease Inhibitor (Roche)) and sonicated (Diagenode Bioruptor, Seraing, Belgium) to around 500 bp DNA fragments. Debris were removed by centrifugation and the lysate was diluted 1:4 with IP dilution buffer (20 mM Tris-HCl pH 8.0, 0.15 M NaCl, 2 mM EDTA, 1% TX-100, protease inhibitors) and precleared with Affi-Prep Protein A support beads (Bio-Rad). The respective antibodies were incubated with the lysate overnight at 4°C, followed by 2 hours incubation at 4°C with blocked protein A Affiprep beads (Bio-Rad) (blocking solution: 0.1 mg/ml BSA). Beads were washed with washing buffer I (20 mM Tris-HCl pH 8.0, 0.15 M NaCl, 2 mM EDTA, 1% TX-100, 0.1% SDS, 1 mM PMSF), washing buffer II (20 mM Tris-HCl pH 8.0, 0.5 M NaCl, 2 mM EDTA, 1% TX-100, 0.1% SDS, 1 mM PMSF), washing buffer III (10 mM Tris-HCl pH 8.0, 0.25 M LiCl, 1 mM EDTA, 0.5% NP-40, 0.5% sodium desoxycholate) and TE-buffer (10 mM Tris-HCl pH 8.0,1 mM EDTA). Beads were eluted twice (25 mM Tris-HCl pH 7.5, 5 mM EDTA, 0.5% SDS) for 20 minutes at 65°C. The eluates were treated with proteinase K and RNase for 1 hour at 37°C and decrosslinked at 65°C overnight. The samples were further purified by phenol-chloroform extraction and ethanol-precipitated. The pellet was dissolved in TE buffer.

For NIPBL ChIP-seq experiments two different protocols were applied. Initially, we used a previously published protocol employing only formaldehyde crosslinking (FA-xlink)^6^. Subsequently, we employed a protocol (DSG/FA-xlink) adapted from van den Berg et al.^43^ that has been shown to allow efficient detection of weaker NIPBL binding sites. For this second protocol, before crosslinking with formaldehyde, cells underwent a protein-protein crosslinking step. In brief, cells were suspended in PBS and treaded for 45 minutes at room temperature under rotation with 2 mM disuccinimidyl glutarate (DSG). After three washes with PBS, cells were crosslinked with formaldehyde as described above. Sonication and pull-down were performed as described above with modifications of the beads washing steps according to Kagey et al.^44^. Beads were washed once with the IP dilution buffer, once with 20mM Tris-HCl pH8, 500mM NaCl, 2mM EDTA, 0.1% SDS, 1%Triton X-100, once with 10mM Tris-HCl pH8, 250nM LiCl, 2mM EDTA, 1% NP40 and once with TE buffer containing 50 mM NaCl.

For sequencing, the DNA libraries were prepared using the NEXTFlex ChIP-Seq kit (BioO Scientific, Austin, TX, USA). These libraries were sequenced according to the Illumina TruSeq v3 protocol on an Illumina HiSeq2500 sequencer (Illumina, San Diego, CA, USA). Single reads were generated of 50 base pairs in length. The quality of DNA sequence was investigated using FASTQC (version 0.11.2), and, when necessary trimmomatic (version 0.32) was used to remove low-quality reads and regions. Quality controlled sequence was aligned to Human genome (hg19) using bowtie (version 1.0.0), and samtools (version 0.1.19) was used to remove reads with mapping quality less than 30, and to keep only aligned reads. Duplicated reads were removed, after alignment, using Picard (version 1.97). Peaks were called using MACS (macs 2) and Peaks were filtered using P-value. UCSC tracks were generated after duplicate removal, using HOMER (version 4.3) and deeptools (version 3.0.2). Heatmaps were generated using Deeptools (version 3.0.2) and NGS plot (version 2.61). Peaks were annotated to specific regions using ChIPseeker (Bioconductor package version 1.14.2).

### ChIP-qPCR

For ChIP-qPCR experiments the ChIP for IgG control, SMC3 and NIPBL (NIPBL #1 antibodies) were performed as described above for the SMC3 ChIP-seq experiment. qPCR analysis using Platinium taq (Thermo Fisher Scientific) was performed according to the manufacturer’s instructions and analysed using a CFX96 C1000 Thermal cycler (Bio-Rad) using the qPCR primers listed in Supplementary Table 1.

### Precocious sister chromatid separation assays

Cells were treated for 30 minutes with 100 ng/ml of nocodazole. Cells were collected and resuspended in 1 ml of medium. 1.5 ml of tap water was added. 6 minutes later 7 ml of Carnoy fixative (3:1, methanol: glacial acetic acid). Cells were then spread on glass slide, dried, stained with Giemsa stain and mounted in Entellan. Mitotic chromosome spreads were observed using a light microscope (DM2000 Leica, France) with a 40x dry objective.

### Fluorescence *in situ* hybridisation (FISH)

DNA FISH was performed as previously described^45^, with the exception that cells were fixed as described above and processed for FISH after spreading on glass slides. Only pairs for which the dots could be clearly resolved were considered in the analysis. Microscopic image acquisitions were performed on a Zeiss Axio Imager M2 using 63X oil-immersion objective. Image analysis was performed using ImageJ software.

### Whole Genome Sequencing

Genomic DNA isolation was performed from blood with the DNeasy Blood & Tissue Kits (Qiagen, Hilden, Germany) according to the manufacturer’s instructions. Genomic DNA quantity was assessed using the Qubit dsDNA BR Assay Kit (Thermo Fisher Scientific). For library preparation, 1000 ng of genomic DNA were used together with the TruSeq DNA PCR-Free Kit (Illumina) following the manufacturer’s recommended protocol. The genomic DNA was fragmented to an average length of 350 bp by sonication on a Covaris E220 instrument (Covaris Inc., Woburn, MA, USA). Library preparation was performed in an automated manner using the NGS Option B (Agilent Technologies, Santa Clara, CA, USA). We assessed the library quality and absence of primer dimers by running a Bioanalyzer DNA High Sensitivity chip. Library quantification was performed using qPCR together with the Kapa Library Quantification Illumina/Light Cycler 480 (Roche). The validated libraries were pooled in equimolar quantities and sequenced via 150 bp paired-end on an Illumina HiSeq 4000 platform following Illumina’s recommended protocol. Raw data were demultiplexed with the Illumina bcl2fastq 2.17 into individual fastq files. Reads were aligned using the mem algorithm of bwa 0.7.8 and aligned to the hg19 reference into which decoy sequences had been added, PAR regions had been masked and the mitochondrial DNA had been replaced with the rCRS (revised Cambridge Reference Sequence) to match MITOMAP and most publicly available resources for mtDNA variants. Base quality scores were recalibrated using GATK BaseRecalibrator with enlarged context size for SNVs and Indels with respect to default (4 and 8 base pairs instead of 2 and 3). Variant calling was then performed using first HaplotypeCaller, to produce an individual GVCF file, and then multisample calling was performed using CombineGVCFs and then GenotypeGVCFs which produced the actual variant calls. Variant qualities and filters were then assessed with VariantRecalibrator tool using the tracks from GATK bundle and variants from GnomAD. For *de novo* variants, the Genotype Refinement workflow was applied using GenotypePosteriors to which a PED file with the family relationships, and again, a GnomAD track with allele counts and frequencies were supplied. The two subsequent steps were VariantFiltration to exclude low quality genotypes and VariantAnnotator to annotate possible *de novo* in the final VCF file.

### Mammalian two-hybrid assay

A fragment of NIPBL containing amino acids 1–300 was inserted into the pCMV-BD expression plasmid (#211342, Agilent Technologies, Santa Clara, CA, USA). The full-length open reading frame of MAU2 was cloned into the pCMV-AD plasmid (#211343, Agilent Technologies). MAU2 mutant constructs containing the deletion identified in the patient of our cohort (c.927_947del21; p.(Gln310_Ala316del)) were generated by site-directed *in vitro* mutagenesis with the Quick Change Site-directed mutagenesis kit, according to the manufacturer’s instructions (Agilent Technologies). HEK293 cells were transiently transfected in 24-well plates with FuGene-HD (Promega), according to the manufacturer’s instructions. Each well was transfected with 250 ng of the pCMV-BD-NIPBL1-300aa, 250 ng of the pCMV-MAU2 wild type or mutant constructs, 250 ng of the Firefly Luciferase reporter plasmid (Promega) and 2,5 ng of the phRG-TK Renilla luciferase expression plasmid (Promega). Following incubations of 24 h, expression of fusion proteins was verified by immunoblot analysis. Activity of Firefly and Renilla luciferases was measured with the Dual Luciferase Reporter Assay System (Promega) with the Tristar^2^Multimode Reader LB942 (Berthold Technologies). All measurements were performed in triplicate in at least three independent experiments. Relative luciferase activity, indicating the strength of the interaction, was determined as the triplicate average of the ratio between the Firefly and the Renilla luciferase activity.

### Yeast two-hybrid assay

The first 300 amino acids of NIPBL and the wild type and mutant full-length MAU2 were cloned into the Matchmaker GAL4 Two-Hybrid System 3 (Clontech-Takara) pGBKT7 and pGADT7 plasmids, respectively, to obtain NIPBL–GAL4 BD or MAU2–GAL4 AD fusion proteins. Yeasts (AH109) were co-transformed with the NIPBL and MAU2 fusion proteins, according to the Matchmaker 3 manual. Proper expression was verified by Immunoblot analysis. Growth selection assays were performed using SD agar plates lacking Trp, Ade, His and Leu to detect interacting transformants.

---

**Supplementry Figure 1.**
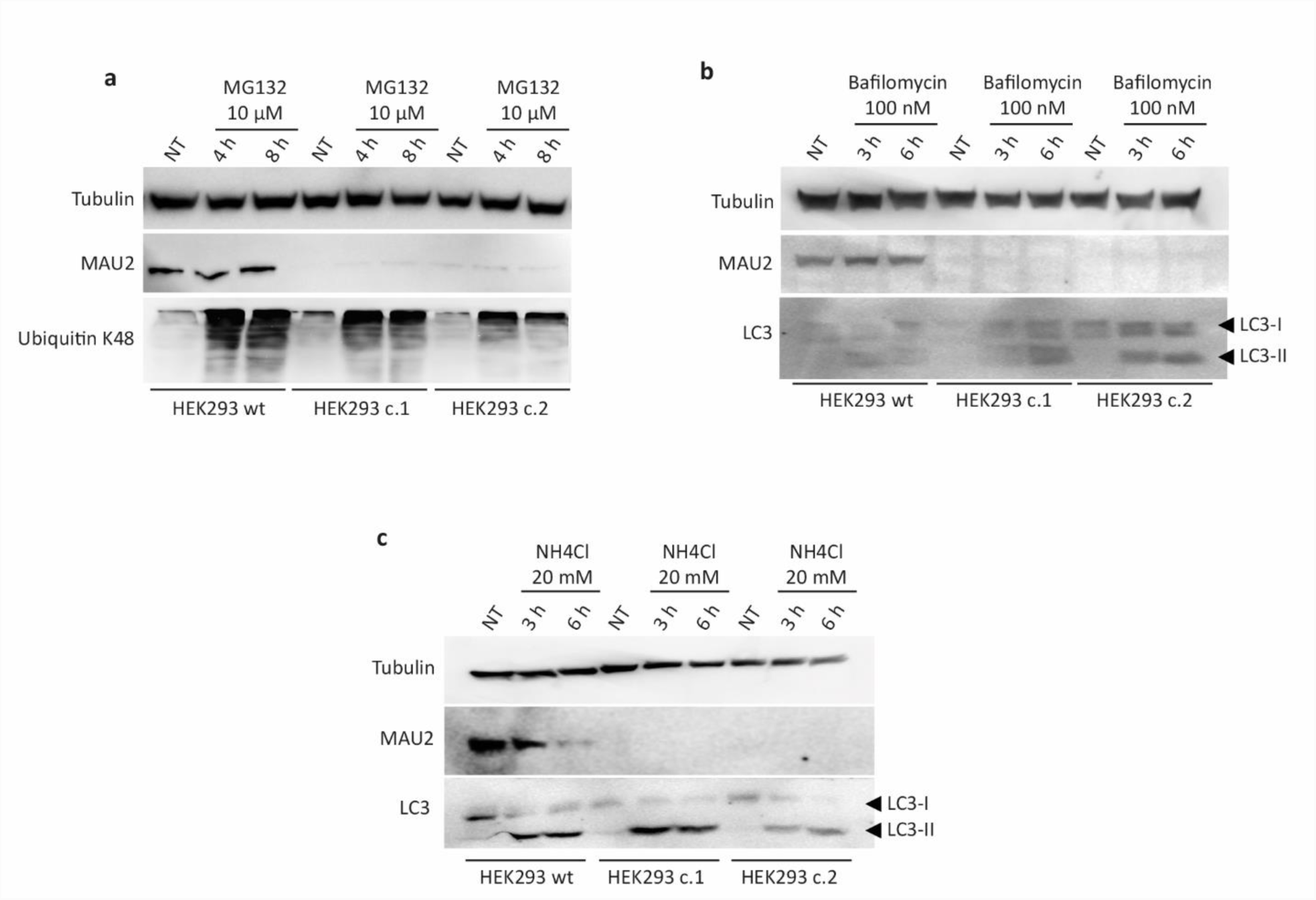

**Supplementry Figure 2.**
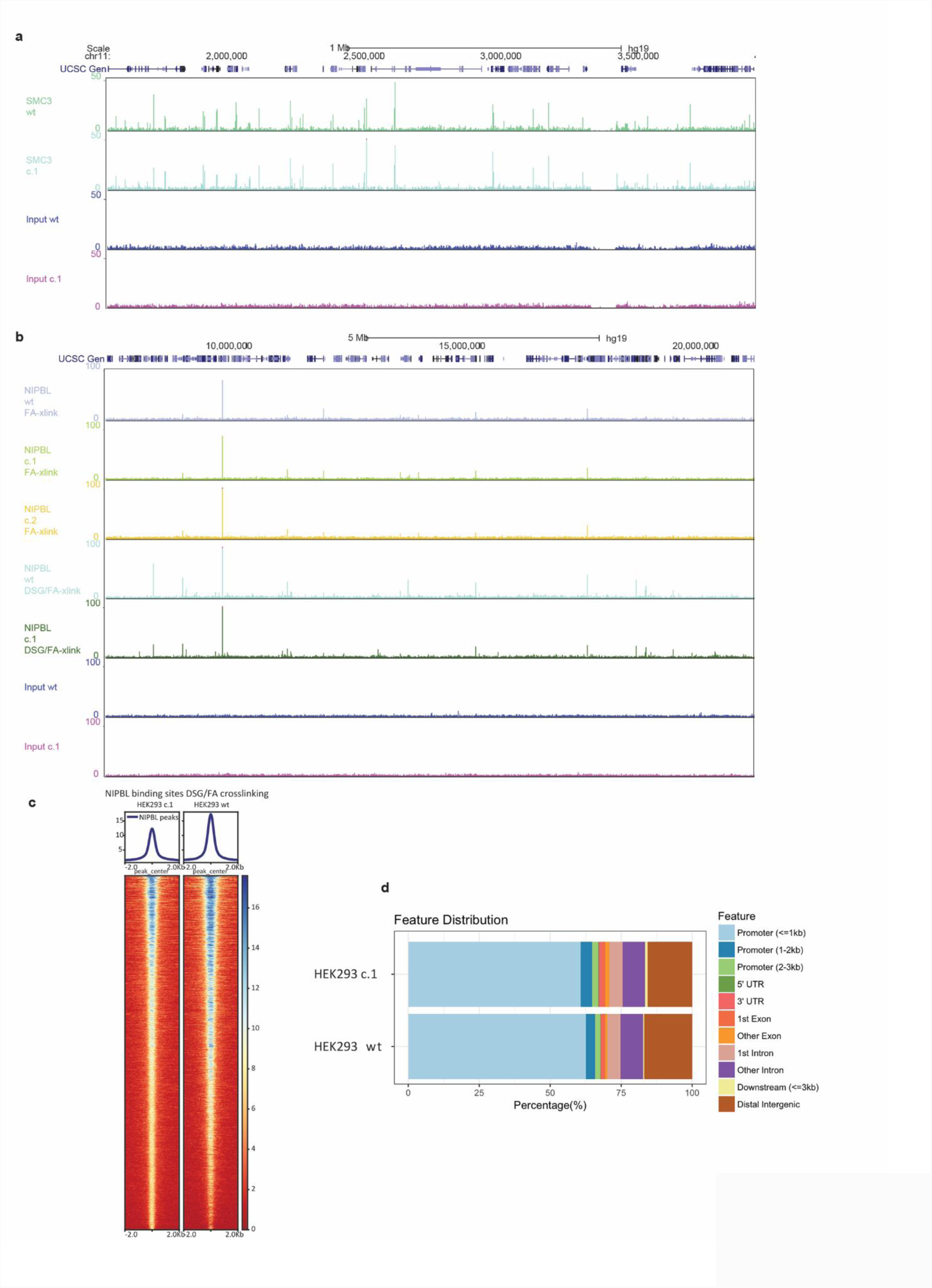

**Supplementry Figure 3.**
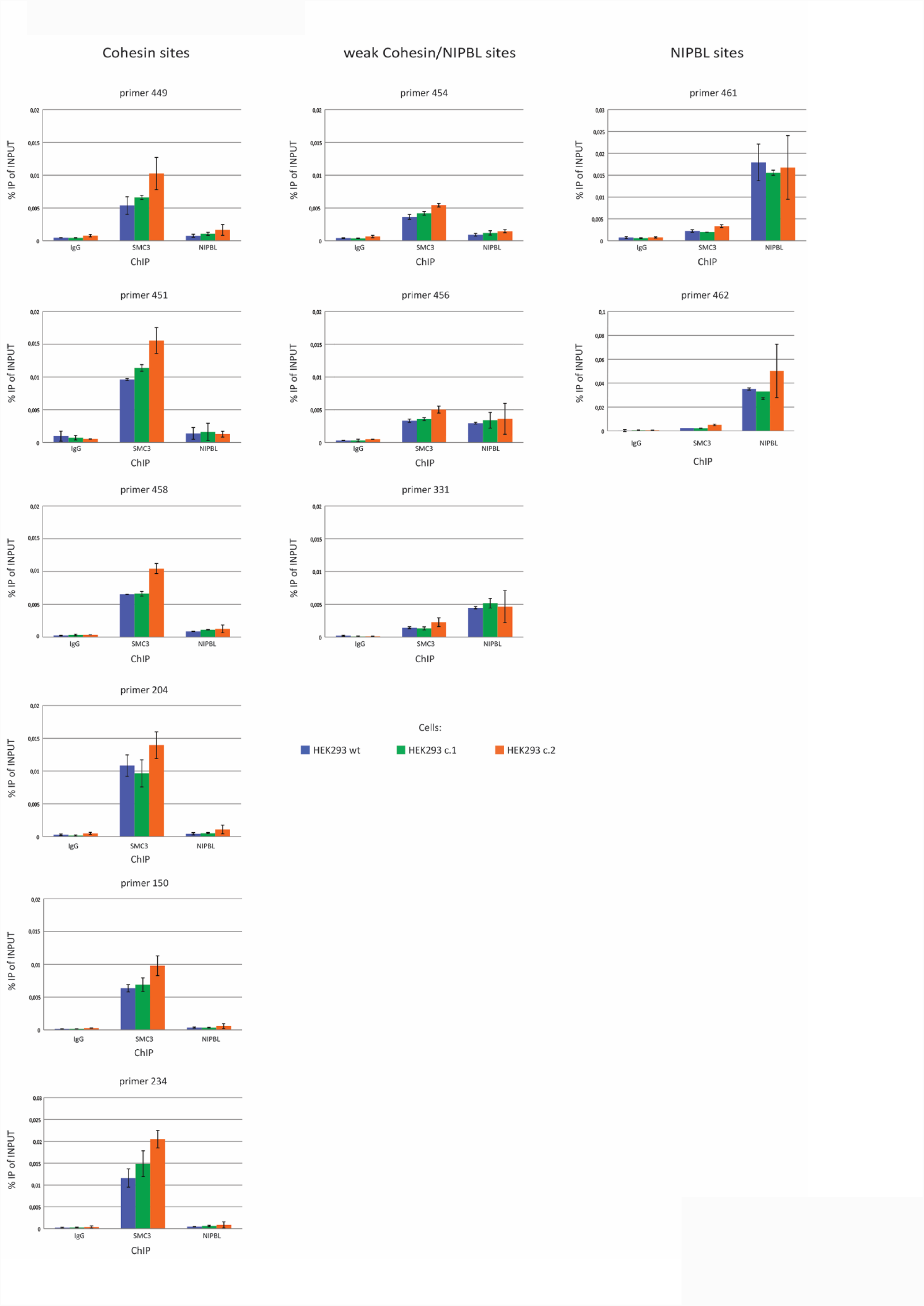

**Supplementry Figure 4.**
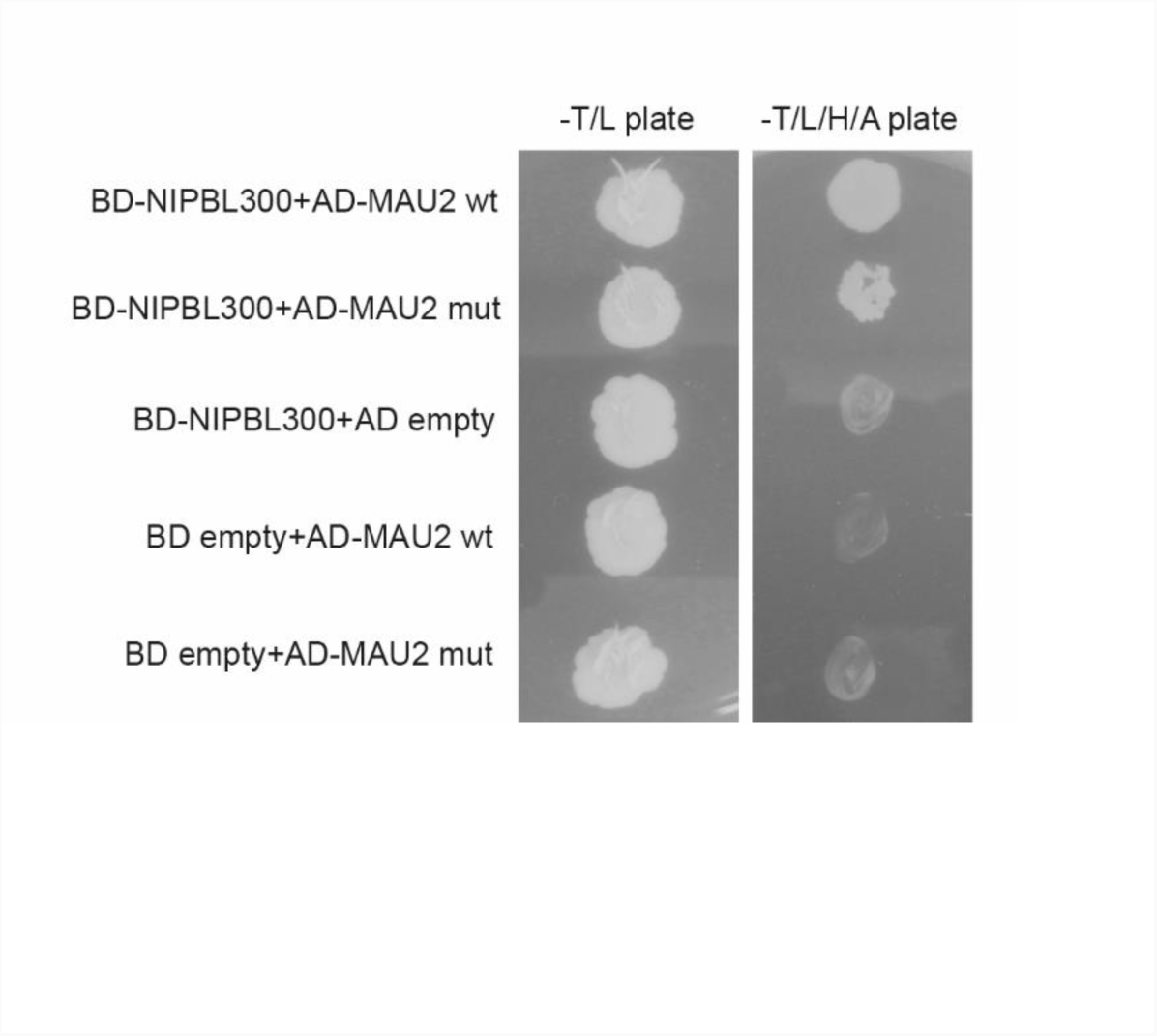

**Supplementary Table 1.**
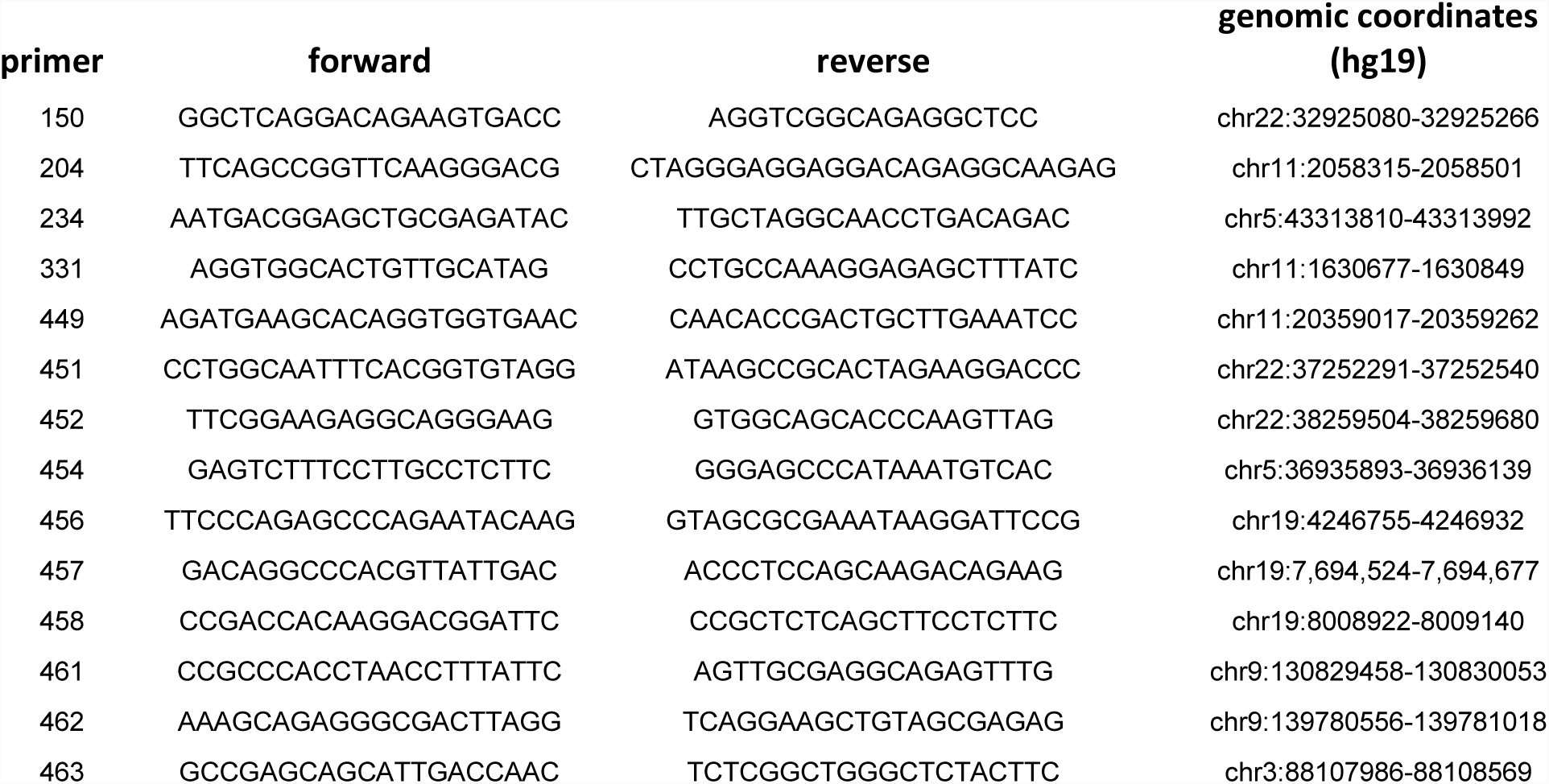

